# Localization of hyphal growth associated with mycotoxin production during the malting of Fusarium head blight infected grains

**DOI:** 10.1101/2020.06.06.126979

**Authors:** Zhao Jin, Shyam Solanki, Gazala Ameen, Thomas Gross, Roshan Sharma Poudel, Pawel Borowicz, Robert S. Brueggeman, Paul Schwarz

**Affiliations:** Department of Plant Sciences, North Dakota State University, Fargo, ND 58108, USA; Department of Crop and Soil Sciences, Washington State University, Pullman, WA 99164, USA; Department of Plant Pathology, North Dakota State University, Fargo, ND 58108, USA; Department of Animal Sciences, North Dakota State University, Fargo, ND 58108, USA

**Author notes:** Corresponding Authors: Shyam Solanki and Paul Schwarz.

**Keywords:** Fusarium head blight, malting, mycotoxins, hyphae, metagenomics, microscopy

## Abstract

Fusarium head blight (FHB) and the occurrence of mycotoxins is the largest food safety threat to malting and brewing grains. Objectives of the current study were to localize the growth of *Fusarium* within FHB infected kernels and to associate it with the production of DON that occurred during malting. FHB infected barley, wheat, rye, and triticale grains that exhibited large increases in *Fusarium Tri5* DNA and trichothecene mycotoxins following malting, were screened for hyphal localization. The growth of hyphae, both on the surface of kernels and within tissues of grain and malt was, imagined by scanning electron microscopy and confocal laser scanning microscopy assisted with WGA-Alexa Fluor 488 pre-staining, respectively. In barley, hyphae were primarily present on or within husk, vascular bundle, and pericarp cavities. Following malting, large amounts of hyphal growth were observed in not only these regions, but also in the aleurone layer, endosperm, and embryo. Extensive fungal growth was also observed following malting of wheat, rye, and triticale. Interestingly, these grains already had an extensive internal presence of hyphae in unmalted grain, occurring in the pericarp, testa, vascular bundle, nucellar projection, aleurone layer, endosperm, pericarp and endosperm cavities, and embryo. Shotgun sequencing followed by metagenomics analysis verified that *Fusarium* spp. accounted for above 90% of the fungal hyphae growing in the interior of grains during malting, which coincided with the significant production of mycotoxins.

## INTRODUCTION

The occurrence of mycotoxins is the largest food safety threat to raw materials in the malting and brewing industries, and recent worldwide surveys of commercial beers have shown deoxynivalenol (DON) to be a frequent, albeit, low-level contaminant (Kostelanska et al. 2009; Varga et al. 2013; Vidal et al. 2016; Wu et al. 2017; Peters et al. 2017). The source of mycotoxins is brewing grains including malt and cereal adjuncts, and in several cases, wheat beers have been reported to have higher DON levels than conventional barley-based beers (Scott 1996; Bauer et al. 2016). The production of trichothecenes, including DON, on kernels of small grains is associated with Fusarium Head Blight (FHB) or scab. In North America, infection is primarily caused by *Fusarium graminearum*, although other species are involved (Jones and Mirocha 1999; Stack 1999; Puri and Zhong 2010).

Several studies have shown that the DON present on malt is largely extracted into the resultant beer (Lancova et al. 2008; Schwarz et al. 1995). However, as the conditions of germination in malting provide a favorable environment for fungal growth (high grain moisture, high relative humidity, and moderate temperatures), and the production of DON and other mycotoxins, the malting process itself is of concern. The behavior of *Fusarium* spp and fate of DON during the malting of barley has been extensively investigated over the past two decades and was recently reviewed by Schwarz (2017). Results have generally suggested that grains with lower DON levels can often be used in malting as most of the DON present on barley is lost during the cleaning and steeping steps. It has also been observed that the viability of *Fusarium* on barley decreases with time in storage. As such, problems with *Fusarium* growth and production of additional DON during the germination phase can often be reduced, if not eliminated, when infected grain with low DON is not malted in several months after harvest.

In the USA, self-imposed DON limits were enacted by industry to accept barley for malting in the range from 0.5 to 1.0 mg /kg. In this range DON levels typically decrease to a safe level following malting, however, maltsters occasionally observe samples that exhibit aberrant behavior. In these cases, DON levels increase significantly during malting even after the barley has been stored for long periods. Thus, companies still must impose limits of DON in malt. Several maltsters have speculated that the increased DON production during malting of these aberrant samples is a result of “internal *Fusarium* infection”, as opposed to the normal “external infection”. In support of this is the observation that pearling (abrasive milling for removal of outer tissues) of samples exhibiting this behavior does not result in the complete removal of DON which is typically observed (P. Bolin, *personal communication*). Speculation on “external vs internal” infection may have some basis in fact as barley spikes are normally enclosed in the flag leaf sheath during anthesis (Langevin et al. 2004; Alqudah and Schnurbusch 2017), and exposure to the pathogen would occur only after the emergence of spikes. On the contrary, in wheat, rye, and triticale, anthers are fully extruded at flowering, which occurs after spike emergence. In addition, rye is open-pollinated and flowers for an extended period of time (Bonnett 1966). Exposure to the pathogen is earlier for these grains and could result in more extensive colonization within grains, when compared to the later infection of barley.

Recent work by Jin and coworkers somewhat supported this hypothesis (Jin et al. 2018a, 2018b). Their investigation showed dramatic increases in DON following the malting of FHB infected wheat, rye, and triticale. DON levels increased as much as 10-fold following the malting of 137 wheat, rye and triticale samples. While 75% of these grain samples had DON levels below the Food and Drug Administration (FDA) advisory level of 1.0 mg/kg, over 80% of these samples had DON in excess of 1.0 mg/kg following malting. DON levels increased up to 20 mg/kg in some of malted rye and triticale samples that originally had low DON. Moreover, relatively high levels of other type B trichothecenes including 3-Acetyl DON (3-ADON), 15-Acetyl DON (15-ADON), and nivalenol (NIV) were also observed in some of these malts. The growth of toxigenic *Fusarium* was measured by quantifying Tri5 DNA in grains and malts, and levels exhibited a moderately strong relationship with malt DON.

Growth of *Fusarium* in grains during malting is, in a sense, an extension of the infection and colonization that have occurred in the field. Previous studies have suggested that the *Fusarium* infection on grains was linked to the host anthesis and infection timing and patterns (Path and Strange 1971; Prom et al. 1999; Trail 2009). During wheat anthesis, infection occurs when ascospores or macroconidia of *F. graminearum* deposit on or inside flowering spikelets. Hyphae then penetrate the ovary and eventually infect the floral bracts including the paleae, lemma, and glume through stomates. Wheat spikes are susceptible to infection from anthesis through the soft dough stage, and the fungus spreads through the rachis (Pritsch et al. 2000; Wanjiru et al. 2002). While wheat, on average, has three flowers per rachis node, rye has only one flower per node, which results in a less dense head. Lower head density provides for better aeration and additionally presents a longer distance for the fungus to travel when spreading through the rachis. These factors could possibly be associated with the lower accumulation of DON that has been observed for rye and triticale grains, when compared to wheat (Gaikpa et al. 2019). Rye has been reported to have some resistance to FHB (Langevin et al. 2004). However, in barley, where spikes are still enclosed in the flag leaf sheath during anthesis, it has been reported that fungal growth occurs across the surface tissue of florets, prior to entry into natural cervices between the paleae and lemma (Lewandowski et al. 2006).

Interaction of the pathogen with surface tissues of the barley grain, and the differences in heading and flowering times have suggested that florets and their morphological characteristics potentially are determinants of the fungal infection and colonization process (Alqudah and Schnurbusch 2017; Imboden et al. 2018). Imboden et al. (2018) found that *F. granimearum* infected barley mainly through prickle-type trichomes on the outer surface of paleae by trapping conidia and providing fungal penetration sites, compared with the dome-type trichomes that lack this function. In addition, the floret maturity of barley influenced the disease spread on grain kernels because it was found that infections of more mature florets supported the spread of hyphae into the vascular bundle, whereas younger florets did not show this spread. The accumulation of silica in response to the presence of fungi on paleae suggested a role for silica in pathogen establishment. It has also been reported that *Fusarium* occasionally utilized stomates along the vascular bundle ridges of wheat and barley to enter the host tissue (Bushnell et al. 2003; Imboden et al. 2018). *Fusarium* infection occurring largely on surface tissue of barley florets is supported by the work of Clear et al. (1997) who found that dehulling or pearling of hulless barley grains removed over 90% of the *Fusarium* and about 50% of DON. However, it should be noted, that pearling, which is an abrasive method, is not effective for the removal of pericarp tissue within the furrow creases regions.

Laca et al. (2006) studied the distribution of mesophilic microorganisms, including bacteria and molds present in wheat by controlled debranning, plate incubation, and scanning electron microscopy (SEM) of the resultant fractions. The results showed that microbes were mainly localized in the grain coat tissues, including pericarp and the protein-rich layer below the pericarp. Jackowiak and co-workers (Jackowiak et al. 2005; Packa et al. 2008) examined wheat and triticale inoculated with *F.culmorum*. SEM of the surface of kernels and transverse sections, showed that hyphae were mainly observed in the areas of furrow creases, the brush end, the caryopsis coat (an area between the coat and the aleurone layer), air spaces at the base of furrow creases, and the cavity located in the central endosperm. It was reported that the cavities were formed originally in the caryopsis by repeated nuclear division during the differentiation of cell walls in the developing endosperm (Burton and Fincher 2014). Heneen and Brismar (1987) found a pericarp cavity PClocated at the base of furrow creases, and the endosperm cavity below the pigment strand in wheat, rye, and triticale grains. These cavities appeared occasionally in grains and were usually located near or in the central endosperm. They also observed that microorganisms often appeared in the pericarp cavities and sometimes, as well, in endosperm cavities of shriveled triticale grains.

Over the years, fluorescence microscopy has been combined with fungal staining techniques to enhance the visualization of hyphae in host plant tissues, such as leaves and seedlings (Moldenhauer et al., 2008, Duckett & Read, 1991, Hoch et al., 2005, Knight and Sutherland., 2011). These studies demonstrated different fluorescent dyes that can distinguish fungi from plant tissues. However, some flaws, such as the quick fading of fluorescence and binding with plant tissues, have challenges to overcome. Green fluorescent protein (GFP) has also been used to ease the observation of fungal growth in plant tissues. However, the GFP method requires genetically transformed mutants (Rollins et al. 2001) and is not really applicable. Recently, Solanki et al. (2019) developed a sensitive method using WGA-Alexa Fluor 488 fungal staining, combined with examination under a confocal laser scanning microscopy (CLSM). This has enabled the imaging of fungal growth within wheat and barley leaves, and with the Z-stack function of CLSM, three-dimensional images showing the penetration of fungi into host plant tissues are possible.

The objectives of this study were to image fungal hyphae within FHB infected small grains and their malts, and then to determine if there was any association between fungal localization and the amount of mycotoxin production observed during malting. Barley, wheat, rye, and triticale grains were all examined because of differences in flowering, and our previous observations on DON production during malting. CLSM examination combined with WGA-Alexa Fluor 488 fungal staining was adapted from the original method of Solanki et al. (2019). Modifications in the sample preparation were required to overcome challenges for application to grain and malt tissues. SEM was used as a supplementary technique to localize hyphae, in consideration of the fact that the extensive washing steps during pre-staining for CLSM might have removed some nonadherent hyphae from host tissues. Metagenomic analysis was utilized to identify the primary fungal microorganisms within these grains and malts.

## RESULTS

### Mycoflora changes, *Fusarium* growth, and trichothecene production during malting

Pink grains were observed in most malt samples selected in the current study. For example, Figure 1 compares unmalted and malted rye grain. While the rye grains appeared clean, large amounts of pinkish kernels were visible following the malting. Figure 2 compares infected grains of barley, wheat, rye, and triticale selected for this study with noninfected (control) samples that were selected from other grain lots. Pictures of infected malt grains are also presented. *Fusarium* Tri5 DNA and type B trichothecenes were not detectable in the control samples of grains and malt (e.g. Tri5 DNA level < 1.0 ng/kg and DON <0.5 mg/kg). While the kernels selected from FHB infected wheat, rye, and triticale samples had some whitish discoloration and/or were shrunken, the appearance of the infected barley was not much different than the control. However, some malt kernels produced from FHB infected grains displayed a pink color. Levels of *Fusarium* Tri5 DNA and type B trichothecenes were considerably higher in the infected samples, which will be discussed subsequently.

**Figure 1.**
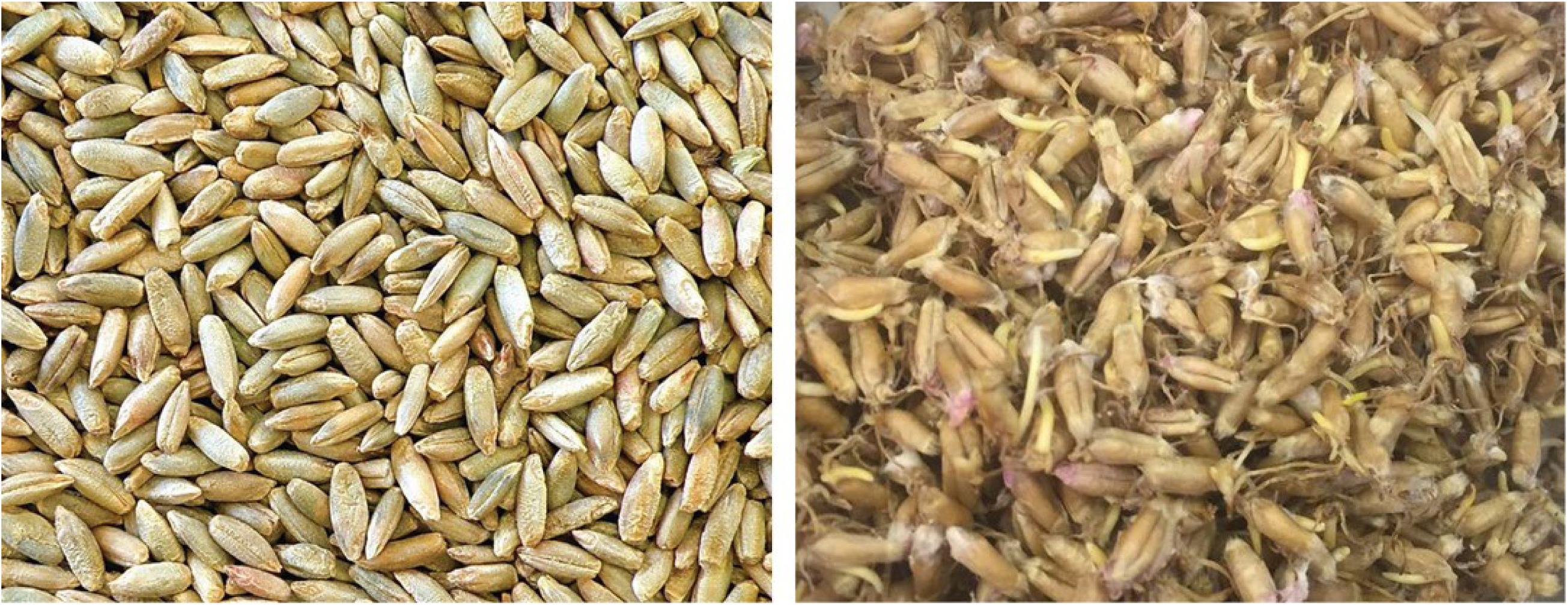
Visible fungal growth on malted rye. A: Rye grain with the DON level below 0.5 mg/kg; B: the corresponding malted rye with the DON level of 9.35 mg/kg.

**Figure 2.**
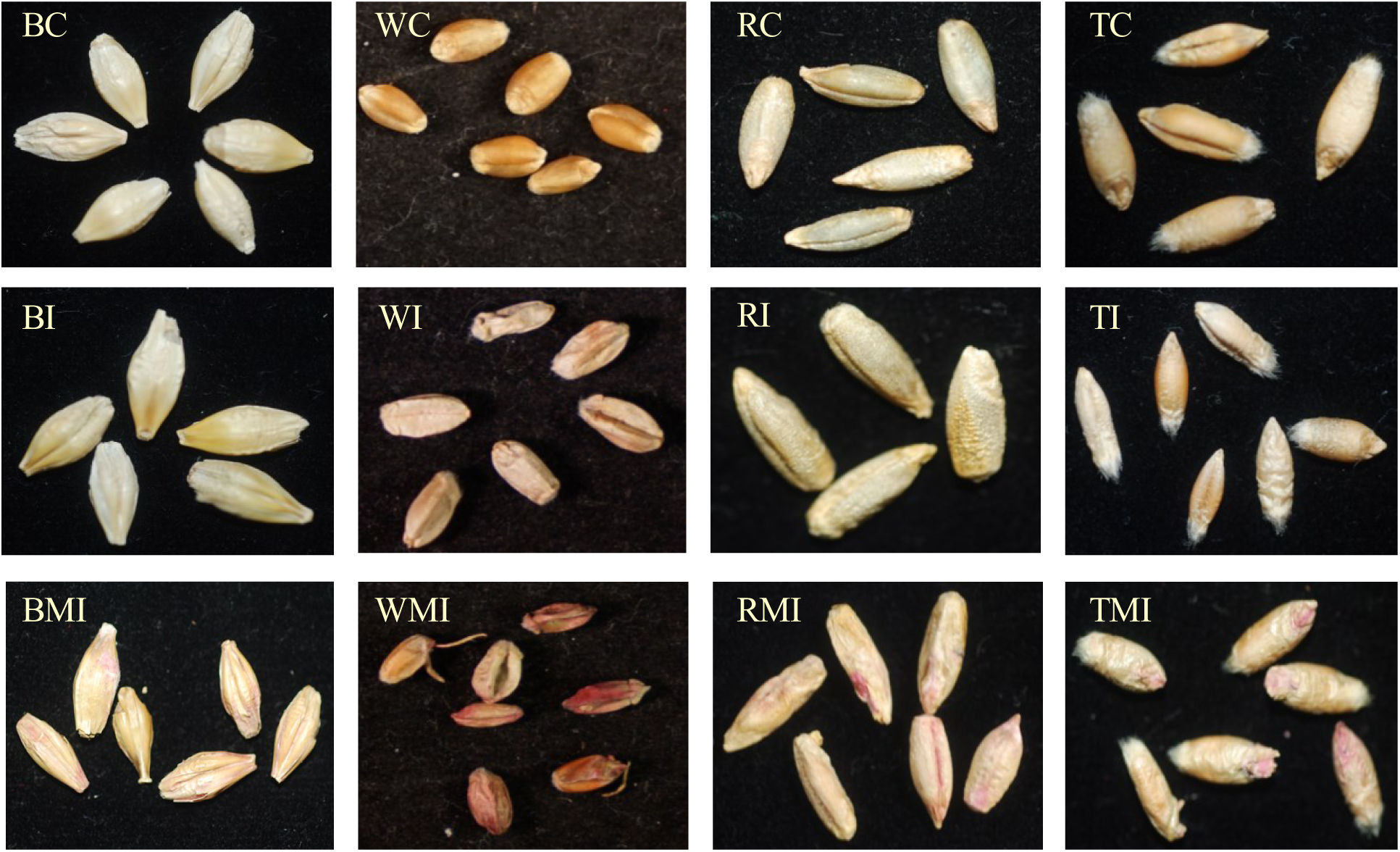
Comparison of healthy and infected grain and malt kernels. BC, WC, RC, and TC are short for barley, wheat, rye, and triticale control samples; BI, WI, RI, and TI are short for barley, wheat, rye, and triticale samples exhibiting FHB infection; BMI, WMI, RMI, and TMI are short for malted barley, wheat, rye, and triticale samples produced from FHB infected grains.

Metagenomic analysis was conducted on all of the grain and malt samples in order to identify microflora. On whole grains, bacteria predominated, and *Fusarium* was actually not even found within the top organisms (data not shown). This was not completely surprising as most grain microflora are known to occur on the outer tissues, and previous work has shown bacteria to be predominant (Priest and Campbell 1996). As our intent was to evaluate internal infection, the results from the outer tissues of whole grains likely masked those of the interior, and it was necessary to remove the husk and pericarp tissues from some grains.

Figure 3 shows the results of mycoflora in dehusked/decoated grain and malt samples, with the analysis of top 270 species detected. Following the malting of wheat, rye, and triticale, the ratio of reads matching *F. graminearum* to the total sequencing reads of DNA extracted from decoated samples increased from 2% to 21%, from 7% to 71%, and from 15% to 47%, respectively. Other *F*. species, including *F. avenaceum, F. oxysporum, F. poae*, and *F. pseudograminearum* accounted for less than 10% of the total reads of DNA extracted from these decoated malt samples. It should be mentioned that the total DNA extracted from decoated samples might not represent the total DNA of host plants and microorganisms because some embryos were probably lost during the decoating process. Generally, the reads matching *F*. species accounted for above 97% of the total fungal microorganisms in each of the decoated wheat, rye, triticale grain, and malt samples.

**Figure 3.**
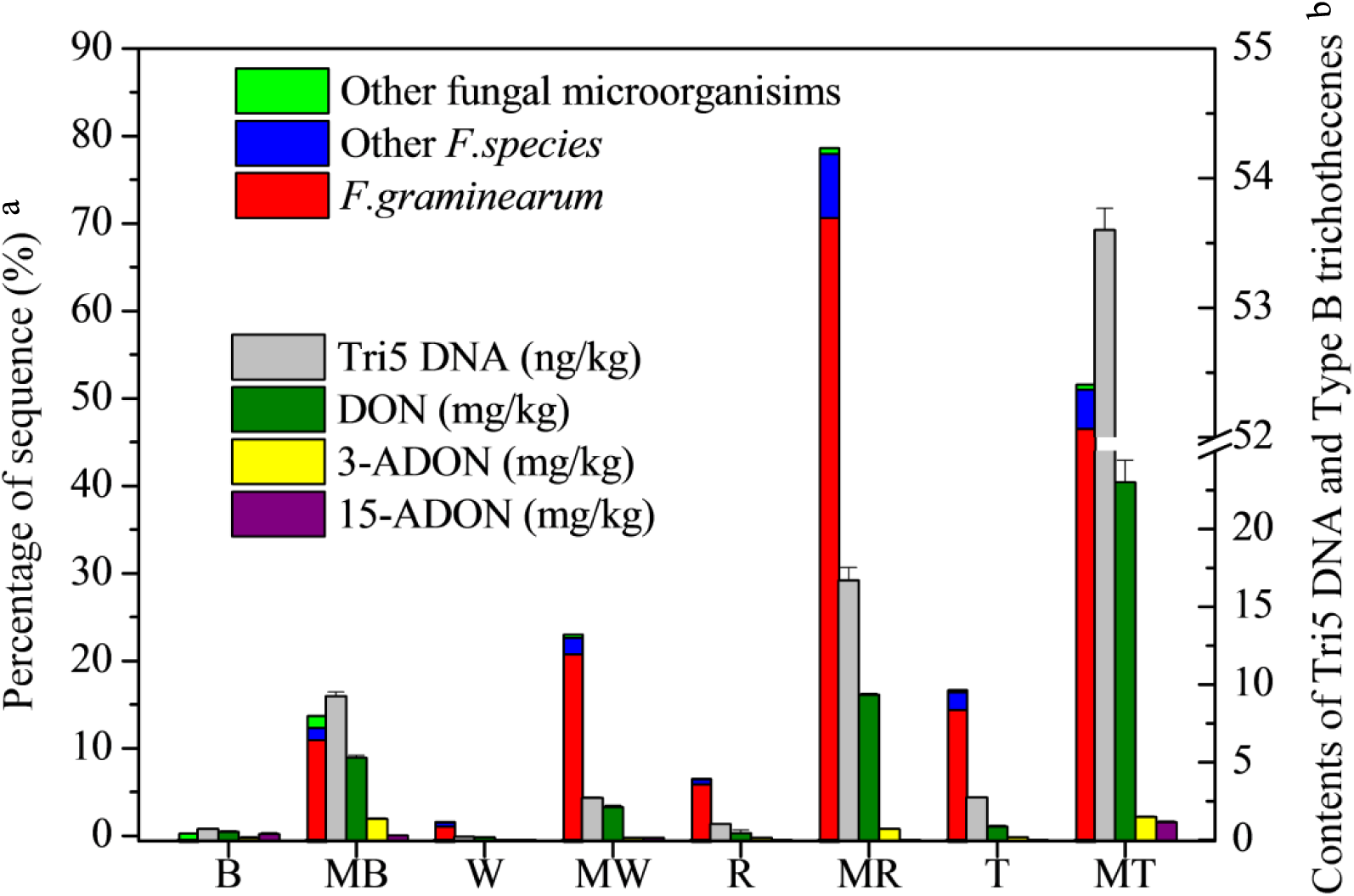
Mycoflora changes, *Fusarium* growth and trichothecene production in the malting of barley, wheat, rye, and triticale. B: Barley; MB: Malted barley; W: Wheat; MW: Malted wheat; R: Rye; MR: Malted rye; T: Triticale; MT: Malted triticale. a: The percentage of DNA reads of *F. graminearum*, other *F.species*, and other fungal microorganisms accounting for the DNA reads of top 270 species detected. The DNA used for metagenomic analysis was extracted from washed dehusked/decoated grain and malt kernels, except for the barley with husk (B). No *Fusarium* detected in washed dehusked barley, thus metagenomics result for this sample is not shown here. b: Quantification of Tri5 DNA and type B trichothecenes were analyzed on whole grain and malt samples.

In dehusked malted barley samples, sequencing reads associated with fungal microorganismsaccounted for 14% of the total DNA reads. *F. graminearum* and other *F*. species constituted 81% and 10% of the total fungal microorganisms, respectively. Fungal genera, including *Alternaria, Aspergillus, Aureobasidium, Malassezia, Penicillium, Rhizopus*, and others totally accounted for less than 10% of the fungal microorganisms in the interior of malted barley. However, *Fusarium* was not found in dehusked barley grains. In addition, mycoflora only accounted for 0.2% of the microorganisms detected in the dehusked barley. Alternatively, with whole barley grains fungal microorganisms accounted for less than 1% of the total sequencing reads, with only 8% of these fungal sequences matching *Fusarium*.

The data presented in Figure 3 also demonstrates that large amounts of trichothecenes were produced during the malting process in the samples of barley, wheat, rye, and triticale. This was accompanied by dramatic increases in *Fusarium* Tri5 DNA levels. DON levels increased from 0.54 to 5.36 mg/kg for the malted barley, from < 0.20 mg/kg to 2.13 mg/kg for the malted wheat, from 0.45 mg/kg to 9.35 mg/kg for the malted rye, and from 0.86 mg/kg to 23.81 mg/kg for the malted triticale. While contents of 3-ADON, and 15-ADON were below 0.50 mg/kg in all the unmalted grain samples, they were produced during malting. The content of 3-ADON increased to above 1.0 mg/kg in the malted triticale and barley, and the 15-ADON content above 1.0 mg/kg in the malted triticale. The level of Tri5 DNA increased by 12.0, 11.9, 15.2, and 18.7-fold in the malted barley, wheat, rye, and triticale, respectively. Lower Tri5 DNA levels in malted barley and wheat samples likely reflected the use of kilned malt, as opposed to the malted rye and triticale, where freeze-dried samples were used.

### Images of fungal hyphae in *Fusarium* infected barley, wheat, rye, and triticale grain and malt kernels

With the FHB infected barley, samples were imaged with both SEM (Figures 4A-D) and CLSM (Figures 4E-H). The overall fungal infection was associated with the husk and furrow creases regions as indicated with the infection spots shown in Figures 4A, and4E. Hyphae were observed on the surface of husk (Figure 4B), in the crevices under husk (Figure 4C), and in the pericarp cavity of barley kernel (Figures 4D) under SEM. Hyphae were also observed with CLSM in the spongy parenchyma of husk located on the furrow entrance (Figure 4F), within the dorsal vein of husk (Figure 4G), and in the vascular bundle (Figure 4H). There were even traces of hyphae localized in the tissues between testa and aleurone layer (Figure 4G), and in the nucellar projection transfer layer (NPTL) (Figure 4H). However few hyphae were found in the aleurone layer, starchy endosperm, and nucellar projection. With malted barley, the extensive distribution of hyphae within kernels is shown in Figures 4I-P. Figure 4I shows the transverse section of a malted barley kernel, and the spots of 4J-P in this figure shows the infection locations observed in multiple kernels of malted barley. With SEM heavy hyphae were observed to fill the furrow creases regions (Figure 4J), and crevices within the embryo (Figures 4K, and 4L). Large amounts of hyphal penetration through grain tissues were observed with CLSM. For insistence, heavy fungal growth was found to occur within the husk, pericarp, testa, aleurone layer, and even endosperm (Figure 4M). Figure 4N shows the hyphae penetrated through the endosperm transfer layer (ETL) and NPTL. The ETL and NPTL structurally refer to the cavity-surrounding cells, as shown in Figure 4N. these layers were found to be responsible for the nutrient transfer from maternal to filial tissues, as such, represent an important role in nourishment and solute supply for endosperm growth (Zheng and Wang 2011). The infection in transfer layers (Figure 4N) was observed to spread down to the starchy endosperm (Figure 4O), and aside to the endosperm (Figure 4P) when checking the conterminal locations of 4N, 4O, and 4P in Figure 4I.

**Figure 4.**
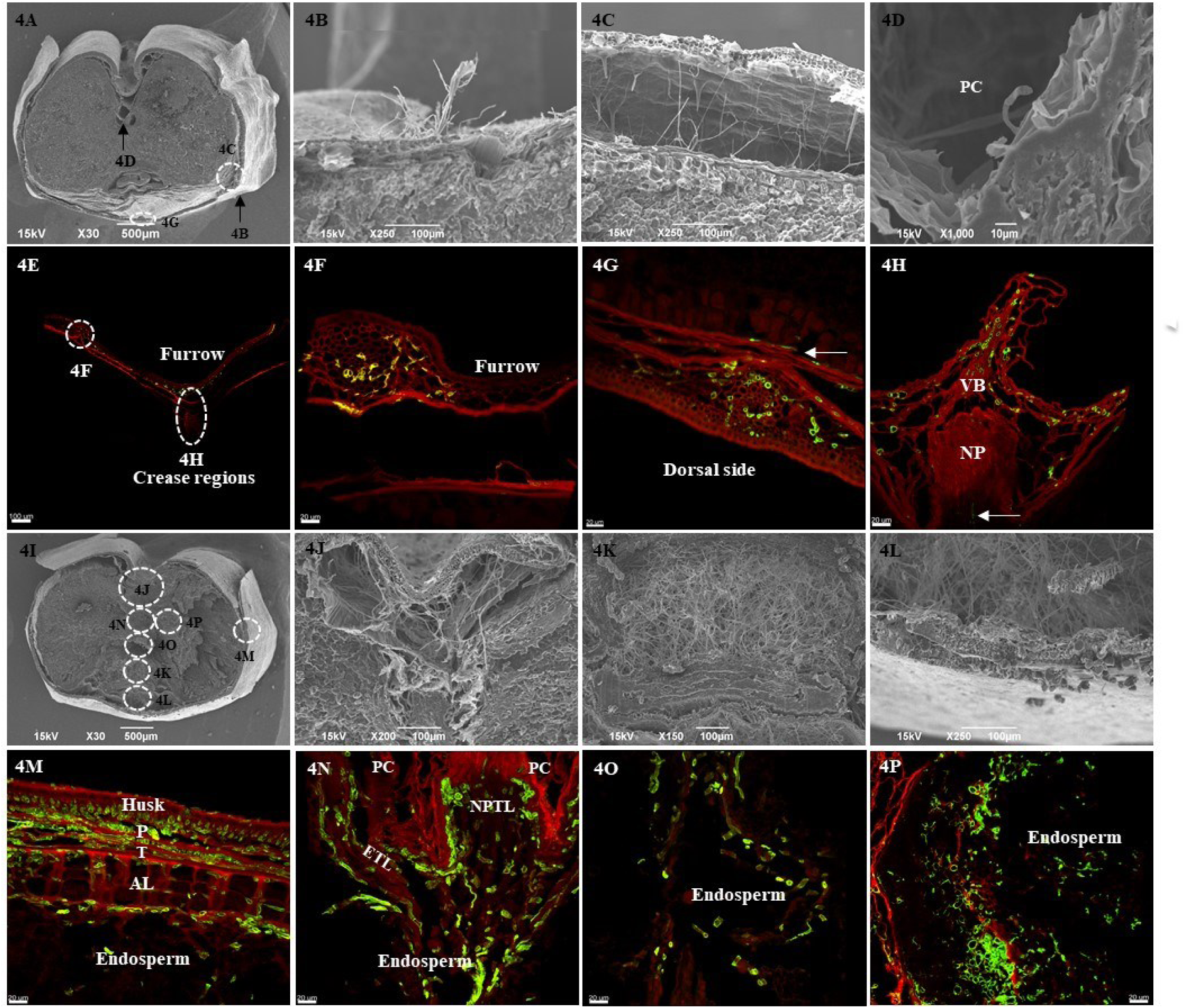
Images of hyphae in the FHB infected barley and malt kernels examined under SEM and CLSM. Figures 4A-D & 4I-L were imaged with SEM, and Figures 4E-J, & 4M-P with CLSM using WGA-Alexa Fluor 488 pre-staining. Figures 4A-H are barley, while Figures 4I-P are malted barley. **A**: Transverse section of an FHB infected barley kernel; **B**: Hyphae on the husk surface; **C**: Hyphae under husk (arrows indicate the prickle-type trichomes); **D**: Hyphae in a pericarp cavity (PC); **E**: Hyphae in the furrow creases (FC) regions; **F**: Hyphae within the spongy parenchyma of husk; **G**: Hyphae in the dorsal vein; Figure **H**: Hyphae within the furrow vascular bundle (VB), nucellar projection (NP), and the arrow indicated a trace of hyphae in the nucellar projection transfer layer (NPTL); **I**: Transverse section of FHB infected barley malt; **J**: Hyphae in the space of PCs, and FC crevices; **K & L**: Hyphae in the crevice of embryo; **M**: Hyphae within husk, pericarp (P), testa (T), aleurone layer (AL), and endosperm; **N**: Hyphae within the cavity-surrounding ETL, and NPTL; **O**: Hyphae within endosperm below Figure 4N; **P**: Hyphae present within the endosperm at the left of furrow.

With the FHB infected wheat, Figures 5A, and 5E show the overall infection spots, which demonstrate the heavy hyphal infection in unmalted kernels. With SEM, hyphae were observed on the surface of kernel pericarp (Figure 5B), and to also extensively within the furrow space (Figure 5C), and pericarp cavities (Figure 5D). With CLSM, hyphae were found to penetrate internal tissues, including the pericarp, testa, aleurone layer, and starchy endosperm (Figure 5F), vascular bundle, nucellar projection (Figure 5G), NPTL, and ETL (Figure 5H). When malted wheat with FHB infection was observed with SEM, there was extensive hyphal growth on the surface of kernel pericarp (Figure 5J), and within the space of cavities and crevices created by modification of the starchy endosperm (Figure 5K, and 5L). Examination with CLSM showed heavy infection in the nucellar projection, NPTL, and ETL (Figure 5N), in the transfer layers, endosperm cavity, and endosperm (Figure 5O), and within the testa, aleurone layer, and endosperm (Figure 5P).

**Figure 5.**
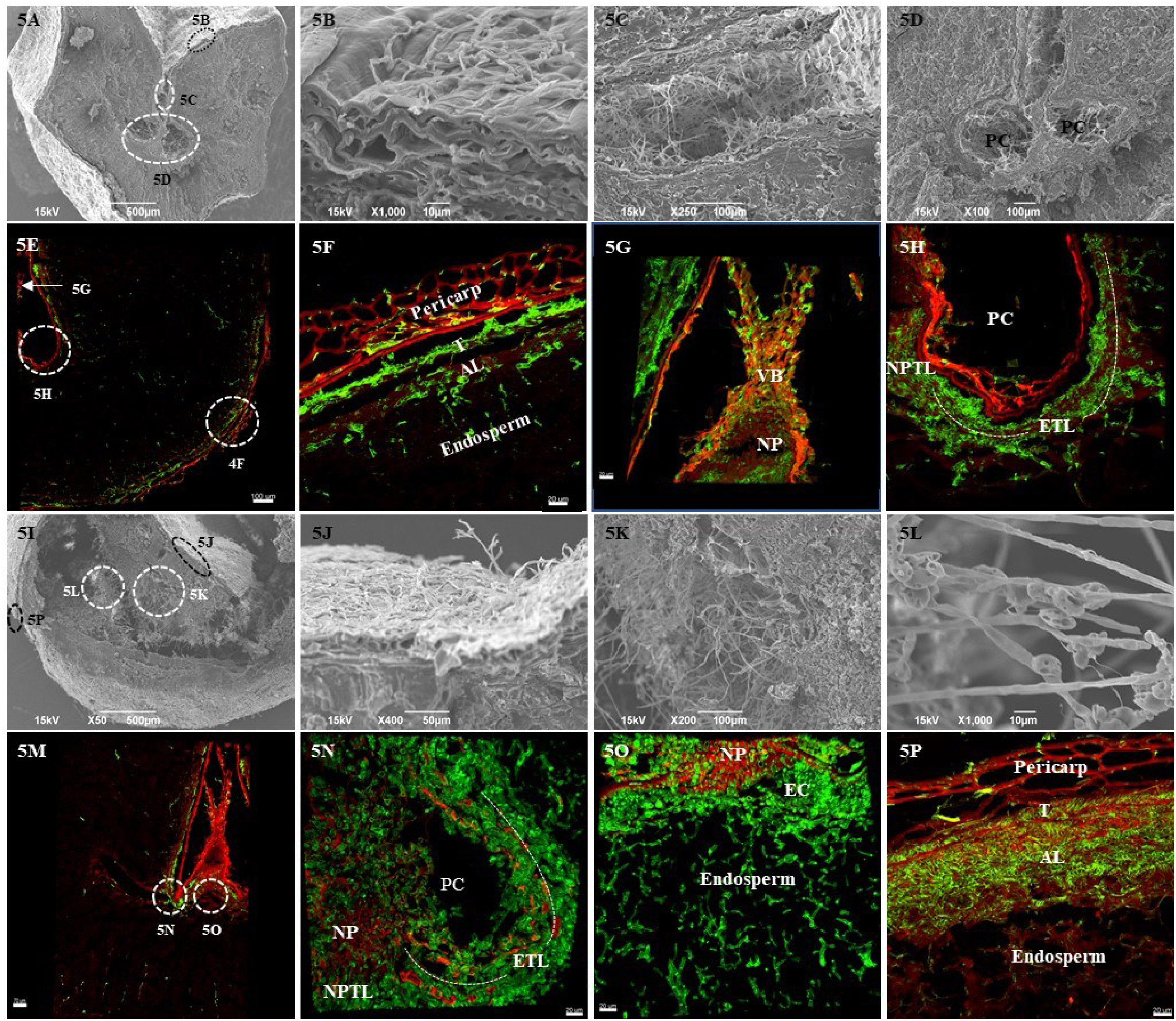
Images of fungal hyphae in the FHB infected wheat and malt kernels visualized with SEM and CLSM. Figures 5A-D & 5I-L were imaged with SEM, and Figures 5E-J & 5M-P with CLSM using WGA-Alexa Fluor 488 pre-staining. Figures 5A-J are wheat, and Figures 5H-P are malted wheat. **A**: Transverse section of an FHB infected wheat kernel; **B**: Hyphae on the surface of pericarp; **C**: Hyphae in the furrow creases (FC); **D**: Hyphae in the pericarp cavities (PCs); **E**: Hyphae in a transverse section of an FHB infected wheat kernel; **F**: Hyphae within pericarp, testa (T), aleurone layer (AL), and endosperm; **G**: Hyphae within the vascular bundle (VB), and nucellar projection (NP); **H**: Hyphae within the NPTL, and ETL; **I**: The transverse section of a FHB infected malted wheat kernel; **J**: Hyphae on the surface of pericarp; **K-L**: Hyphae within crevices of the endosperm; **M**: The transverse section of a FHB infected malted wheat kernel; **N**: Hyphae within the NP, NPTL, and ETL; **O**: Hyphae within the transfer layers, endosperm cavity (EC), and endosperm under EC; **P**: Hyphae within the pericarp, testa (T), AL, and starchy endosperm.

With FHB infected rye and triticale, there were also large amounts of hyphae observed on/within grain kernels. Figures 6A-H show specific spots of hyphal infection present on rye, and malted rye kernels. Hyphae were observed on the surface of testa (from which the pericarp had been peeled off) (Figure 6A), in the space at the base of furrow creases, and pericarp cavities (Figure 6B), between starch granules (Figure 6C), and in the embryo crevices (Figure 6D) under SEM. Figure 6E presents a perspective of general infection with hyphae on a transverse section of a whole rye kernel under CLSM. Hyphae had penetrated into the internal tissues, such as furrow creases tissues, and vascular bundle (Figure 6F), ETL, and endosperm (Figure 6G), pericarp, testa, aleurone layer, and starchy endosperm (Figure 6H). Figures 6I-P shows the hyphal distribution associated with triticale grain and malt. Figures 6I, and 6J show penetration of hyphae through the grain surface into the pericarp. Large amounts of hyphae were observed in the air space of furrow (Figure 6K) and the pericarp cavity (Figure 6L) under SEM. Figure 6M present the overall infection with hyphae on a transverse section of a triticale kernel under CLSM. Hyphae were observed to in the tissues of pericarp, testa, aleurone layer, and endosperm (Figure 6N), furrow creases and the nearby endosperm (Figure 6O), and ETL, NPTL, and endosperm cavity (Figure 6P). The images of FHB infected rye, and triticale malt were not shown because the localization of hyphae was observed to be very similar to that in unmalted grains. However, the hyphal infection tended to be much heavier in malt kernels. Another obvious difference is that the percentage of high DON kernels was found to be much higher in malted rye and triticale samples than that in the corresponding grains, and the specific data would be published in another research article.

**Figure 6.**
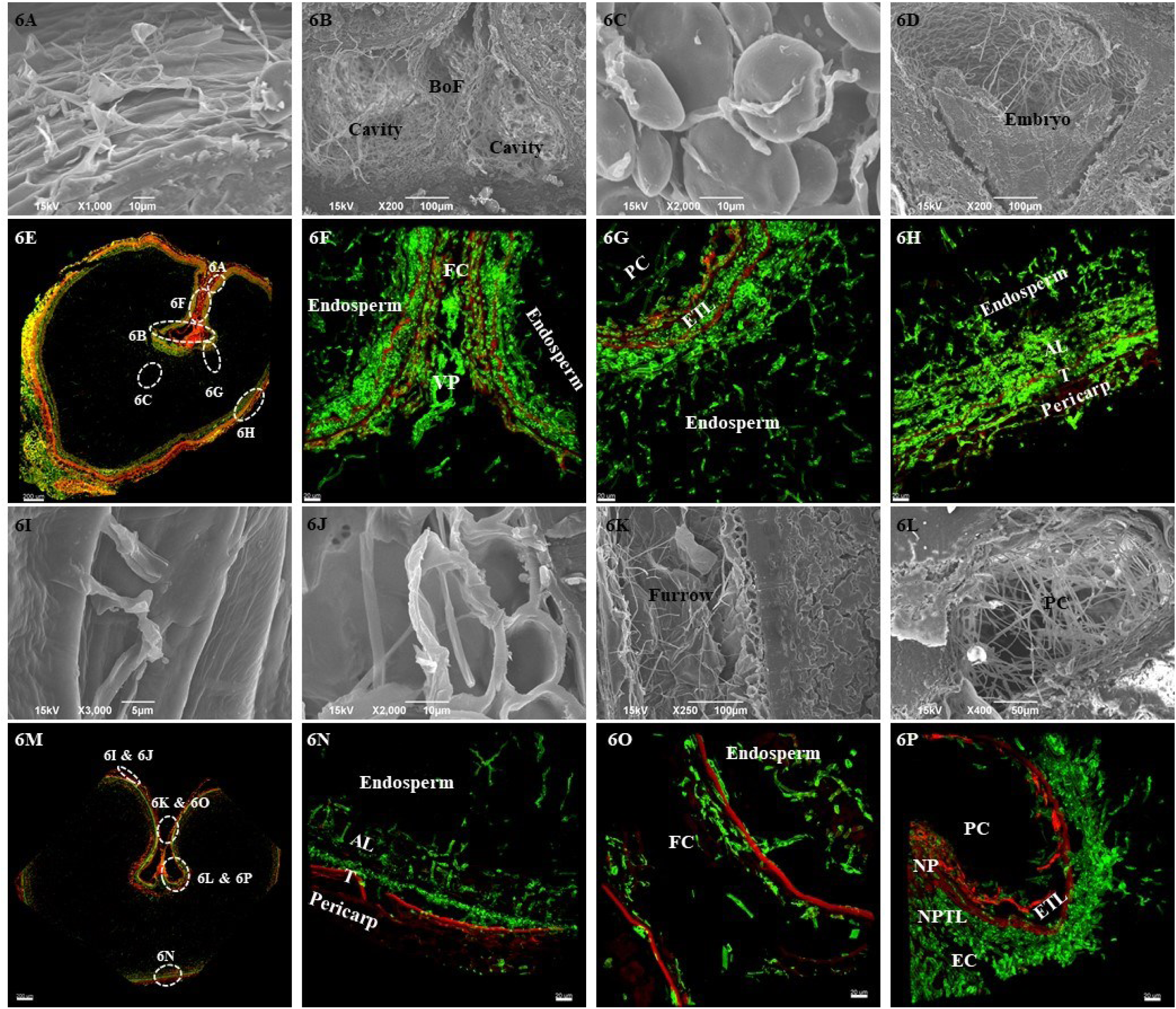
Images of hyphae in FHB infected rye and triticale kernels examined with SEM and CLSM. Figures 6A-D & 6I-L were imaged with SEM, and Figures 6E-J & 6M-P with CLSM using WGA-Alexa Fluor 488 pre-staining. Figures 6A-J are rye, and Figures 6H-P are triticale. **A**: Hyphae on the surface of testa, with the pericarp peeled off; **B**: Hyphae in the air space at the base of furrow creases (BoF) and pericarp cavities (PCs); **C**: Hyphae between starch granules; **D**: Hyphae in the crevices of embryo; **E**: Transverse section of a FHB infected rye kernel; **F**: Hyphae within the furrow creases (FC), VP, and nearby endosperm **G**: Hyphae within the ETL, and endosperm; **H**: Hyphae within pericarp, testa (T), aleurone layer (AL), and endosperm; **I**: Penetration of hyphae through the pericarp surface; **J**: Hyphae in the pericarp; **K**: Hyphae in the air space of furrow; **L**: Hyphae in a pericarp cavity with the location indicated in Figure 6M; **M**: Transverse section of a FHB infected triticale kernel; **N**: Hyphae within pericarp, testa (T), AL, and endosperm; **O**: Hyphae within the FC, and the nearby endosperm; **P**: Hyphae within the nucellar projection (NP), NPTL, ETL, and EC.

Control samples, representing non-infected grain or those with very limited infection, were stained concurrently with those that were FHB infected. As shown in Figure S1, small amounts of hyphae were detected in tissues of these grains and/or malts. In barley, some hyphae were mainly found in the spongy parenchyma of the husk (Figure S1a), and in wheat, within the pericarp of furrow, and sieve element (Figure S1b). Results were similar for rye, with hyphae being detected within the pericarp and testa (Figure S1c), while for triticale hyphae were present in seed coat, and vascular bundle in the furrow region, and even within the ETL under nucellar projection (Figure S1d). However, presence of fungi doesn’t necessarily indicate *Fusarium* infection, as there was no metagenomic data for these control samples.

## DISCUSSION

Maltsters and brewers have for centuries relied on the visual appearance of grain and malt as an indicator of potential microbial problems, and pink grains have been considered to suggest infection with *Fusarium* (Schwarz 2003). Empirically, symptoms such as reddish and black discoloration of kernels (Figures 1, and 2) can be related to *Fusarium* infection, but this symptomatology has generally shown a low predictive value for *Fusarium* contamination because infection by some other fungi like *Rhodotorula* and *Cryptococcus* can cause similar discoloration (Laitila et al. 2011). In the current study, metagenomic analysis demonstrated that *F. graminearum* and other *F*. species were the predominant fungi (>97%) living in the interior of wheat, rye, and triticale, and their malt samples (Figure 3). With barley, *Fusarium* was minimally present in the unmalted grains and mainly distributed in the husk. However, *F. graminearum* grew into the predominant fungus in the interior of barley kernels following malting. This viewpoint was supported by the following evidence. It was observed that only 0.2% of the total reads were identified as fungal microorganisms and *Fusarium* was not even identified in the dehusked grain kernels, whereas *Fusarium* reads accounted for 0.1% of the total DNA extracted from the intact barley grain kernels. The hull was not analyzed in the study, but a previous report indicated that most *Fusarium* was found in the hull fraction pearled from barley (Clear et al. 1997). However, when the husk was removed, *F. graminearum* matched 12% of the total DNA reads from malted barley, and 81% of the fungal DNA reads (Figure 3). The contents of both *Fusarium* Tri5 DNA and DON increased more than ten times following the malting of all the grains, which further demonstrated the dramatic growth of *Fusarium* during malting. This behavior was contrary to a previous report that barley grains with low DON level (e. g. <0.5 mg/kg) produced lower DON on the malt (Schwarz et al. 2006). Differences may be associated with the physical location of *Fusarium* in grain kernels, typically the furrow creases regions, where is not accessible to cleaning and steeping. Imboden et al. (2018) observed that spread of inoculated *F. graminearum* on the furrow vascular bundle tended to occur in mature florets but not in younger ones. This timing of infection and maturity of florets could be possibly influenced by the weather and varieties of barley (Prom et al. 1999). In the earlier study (Schwarz et al. 1995), *F. graminearum* was found in several FHB infected barley samples grown in the upper midwestern United States, and levels of DON and other type B mycotoxins increased following malting. The observation of conidia under the husk surface of germinating barley kernels suggested that the infection would not be completely reduced by the overflow and fill-drain cycles during steeping. In addition, the presence of perithecia and conidia on some germinating barley kernels indicated that the release of conidia could result in the possible cross-contamination.

Imaging using CLSM combined with fluorescent staining was methodologically developed to examine fungal hyphae in barley and wheat leaves (Solanki et al. 2019). In the this method, WGA-Alexa Fluor 488 was used as the fluorescent dye because of its specific binding with N-acetylglucosaminyl residues of chitin in fungal hyphae. However, there were some technical challenges when adopting the method for grain samples which contain large amounts of starch granules. Unlike the transparent leaves, the grain and malt kernels need to be thin-sectioned because the starchy endosperm is nonopaque when scanned with the laser. However, large portions of thin sections of malt kernels were often lost, particularly when malt kernels were significantly modified by germination. This needs to be avoided in order to keep the structure of host tissue intact when sectioned. Solanki et al. (2019) recommended the autoclaving of leaves in a liquid cycle to remove wax from leaf tissues. However, when applied to grain, autoclaving tended to cook the samples and made them too soft to section. As an alternative, we used a series of soaking periods in 10% NBF solution and then varied the thickness of sections. This preliminary work showed that the grain and malt kernels retained their integrity when soaked for 48 hrs before sectioning into 10-15 µm sections. These sections were thicker than the 4-5 µm sections of leaf tissue previously used for microscopic studies (Jääskeläinen et al. 2013; Olkku et al. 2005). In order to improve the permeability of fluorescent stain, we increased the staining time to 5 hrs and replenished the fluorescent solution after 2.5 hrs. The Z-stack images scanned by CLSM had an intact structure of high resolution as shown in Figures 4-6 and S1. Small amounts of hyphae were detected even in the relatively clean grain kernels (Figure S1), indicating that CLSM coupled with WGA-Alexa Fluor 488 staining was a sensitive method to localize hyphae in grain and malt. These hyphae might be inaccessible during cleaning and then become potential sources of hyphal growth in the following malting.

To localize the fungal hyphae within FHB infected grain and malt samples, SEM and CLSM were both used because of their respective advantages. CLSM with the Z-Stack function was capable of imaging hyphae which had penetrated through host tissues, but the extensive washing used in sample preparation, there was a possibility that less adherent hyphae were removed. While SEM is only used to examine the surface of an object, the lack of washing has the advantage of avoiding the loss of hyphae on host tissues. The complementary nature of the two techniques was illustrated by comparing the SEM and CLSM images of the same kernel locations. For instance, hyphae were obviously seen within the cavities of barley and malt (Figures 4D, and 4J), but not visible in the surrounding host tissues when viewed under SEM. In contrast when viewed using CLSM and pre-staining, there was clearly a presence of hyphae within these host tissues (Figures 4H, and 4N). Complementary comparisons of the cavities are also shown in Figures 5D versus 5H for wheat, Figures 6B versus 6F-G for rye, and Figures 6L versus 6P for triticale. SEM examination showed hyphae on the surface of barley (Figure 4B) and under husk (Figure 4C), and on the pericarp of wheat and malt (Figures 5B, and 5J), rye (Figure 6A), and triticale (Figure 6I). In addition, with CLSM hyphae were observed within tissues including spongy parenchyma of husk (Figures 4F, 4G, and 4M), the pericarp of wheat (Figures 5F, and 5P), rye (Figure 6H), and triticale (Figure 6N). Previous studies using SEM confirmed the presence of hyphae on the kernel surface and within the cavities of FHB or shriveled wheat, rye, and triticale grains, and authors speculated on possible infection in the endosperm (Packa et al. 2008; Jackowiak et al. 2005; Heneen and Brismar 1987). In the current study, hyphae were observed physically in crevices of vascular bundle under SEM, and within the tissues and nearby endosperm under CLSM of malted barley (Figures 4J versus 4N-P), wheat (Figures 5C versus 5G), rye (Figures 6B versus 6F-G), and triticale (Figures 6K versus 6O).

In terms of external versus internal infection on kernels, the SEM and CLSM images were aligned to analyze the localization of hyphae on/within each of the grain and malt samples. Only slight infection was observed in the FHB infected barley grain kernels; mostly in the husk and furrow regions of the husk. Hyphae were mainly localized on the husk surface (Figure 4B), in areas under husk (Figure 4C), and in the spongy parenchyma and cementing layer of husk (Figures 4F, and 4G). The distribution of hyphae in the husk likely resulted from the surface infection of *Fusarium* on florets of barley in the field (Langevin et al. 2004), and it is worth noting that the prickle-type trichomes, which have been suggested to have the function of trapping conidia and to act as sites of fungal penetration (Imboden et al. 2018), were observed on the inner surface of barley husk in Figure 4C. Within the furrow, hyphae were present in the vascular bundle (Figures 4E, and 4H) that were created by the fusion of margins of lemma with three layers and paleae with two layers of vascular bundle. Fungal infection at the furrow margin (Figure 4F) was probably caused by the hyphal penetration through stomates that were observed to run along the vascular bundle ridges (Imboden et al. 2018). The furrows and the paleae margins were considered as the frequent location of lesion initiation on grains grown in the field (Lewandowski et al. 2006). In addition, hyphae were observed in cavities under the vascular bundle (Figure 4D), and dorsal vein (Figure 4G). While the infection mechanisms at these two sites were not clear, the inaccessibility of these spots during grain cleaning and steeping prior to germination in malting might be a source of the fungal growth. In a previous study, Olkku et al. (2005) found that DNA stained with DAPI (4’, 6-diamidino-2-phenylindole) was located in pericarp and endosperm of barley and they speculated on the existence of microbes in these tissues. However, DAPI is not specific to microbial DNA. In the current study, extensive growth of hyphae was also observed at these spots following malting of the barley. This resulted in not only the heavy growth of hyphae in the internal crevices like cavities (Figure 4J) and embryo regions (Figures 4K, and 4 L), but also penetration of hyphae into host tissues (Figures 4M-P). The presence of hyphae into the starchy endosperm was facilitated by spread through husk, pericarp, testa, and aleurone layer (Figure 4M), and alternatively through the ETL down (Figure 4N) and aside (Figure 4P) into the endosperm. These results illustrated that even a trace amount of fungi infection in the interior of grains can potentially result in significant growth under favorable conditions of malting.

With FHB infected wheat, rye, and triticale, extensive growth of hyphae were observed both on the kernel surface and within the interior of grains prior to malting (Figures 5A-H, and 6A-H), and infection became even heavier following malting (Figures 5I-P, and 6I-P). The ingrowth of hyphae in the grain tissues, such as aleurone layer and endosperm were not observed in the previous report by SEM. However, Jackowiak et al. (2005) and Packa et al. (2008) both reported that amylolytic enzymes produced by the *Fusarium* used in inoculation were able to diffuse through the aleurone layer and endosperm, causing fractured cells in the aleurone layer and damaged starch granules, which were not even in contact with hyphae. The physical distribution of hyphae was associated with the timing of *Fusarium* infection on these hulless grains, and the pattern of *Fusarium* infection in the field. Wheat, rye, and triticale flower after heading, and the primary infection occurs in the field when ascospores (sexual spores) or macroconidia (asexual spores) are released from soilborne debris and then infect flowering spikelets. Moreover, the infection spreads between florets through the rachis, which results in the fungus entering the base of caryopsis (Gaikpa et al. 2019; Langevin et al. 2004). These patterns of infection could lead to more extensive colonization of *Fusarium* in the interior of wheat, rye, and triticale grain kernels, which provided potential sources of fungal growth and mycotoxin production during malting.

## MATERIALS AND METHODS

### Grain and malt samples

Grains and their corresponding malt samples were from a variety of sources. They were all selected on the basis that they had exhibited large increases in DON following malting, despite the original grain showing relatively low levels. Two-rowed spring barley was from the 2016 commercial crop in Saskatchewan Canada. Both the barley and corresponding malts were provided by a commercial malting company who had identified this as a problematic sample. The hard red spring wheat samples were from the 2015 commercial crop in North Dakota in 2015, and micro-malted in a previous study (Jin et al. 2018b). Rye and triticale samples were from field trials in Minnesota in 2016. Micro-malts of these grains were from a previous study (Jin et al. 2018a). Non- (or slightly FHB) infected barley, wheat, rye, and triticale samples were identified by screening for Tri5 DNA and DON, and used as negative controls for microscopic examination.

### Metagenomic analysis

Metagenomic analysis was performed to determine the identity and levels of fungal organisms associated with the grain and malt samples. However, as the bulk of microbial biomass occurs on the surface of the grain, it can mask changes that occur internally. As such, husk and pericarp tissues were removed from some kernels prior to analysis. For decoating, 1.0 g of each grain or malt sample were washed with 1% of hypochlorous and then soaked in 75% ethanol for 12 hrs. The husk of barley and malt kernels, and the pericarp of wheat, rye, triticale, and the malt kernels were loosened by gently rubbing in a mortar and pestle. Kernels were then peeled with a tweezer to remove the husk or pericarp. The de-husked and de-coated kernels were washed with sterile water and processed with a DNeasy Plant Mini Kit (Qiagen Inc. Valencia, CA, USA) for the fungal DNA extraction. DNA samples were used to generate a whole-genome shotgun metagenomic sequencing library and sequence in Ion Torrent S5 platform. Barcoded-sample specific sublibraries were prepared using NEBNext^®^ Fast DNA Fragmentation & Library Prep Set for Ion Torrent™ kit. The sequencing data were used to perform a blastX search in the regularly updated in house NCBI nonredundant (NCBI-nr) database utilizing the high-throughput alignment program DIAMOND (Buchfink et al. 2015). The microbiome analysis tool MEGAN Community Edition v6.12.3 (Huson et al. 2016) was used to analyze the blastX alignment results generated in DIAMOND and assign individual reads to operational taxonomic unit (OTUs).

### Quantification of *Fusarium* Tri5 DNA

*Fusarium* Tri5 DNA is quantitatively analyzed by qPCR as described by Jin et al. (2018a). The fungal DNA was extracted with a DNeasy Plant Mini Kit (Qiagen Inc. Valencia, CA, USA), amplified with *Fusarium* Tri5 DNA PCR primers TMT_fw (5’-GATTGAGCAGTACAACTTTGG-3’) and TMT_rev (5’-ACCATCCAGTTCTCCATCTG-3’) (Milliporesigma, Burlington, MA, USA), and quantified with SsoAdvanced TM Universal SYBR^®^ Green Supermix (BIO-RAD, Hercules, CA, USA). To quantify the initial amount of Tri5 DNA for each sample, the qPCR reactions are performed simultaneously with a serial dilution of purified *Fusarium graminearum* Tri5 PCR amplicon generated with *F. graminearum* DNA template and extracted from single spore cultures.

### Detection of type B trichothecenes

DON, 3-ADON, 15-ADON, and NIV were determined according to our standard laboratory procedure (Jin et al. 2018b; Wan et al. 2018). Ground samples (2.50 g) was extracted with 20 mL of acetonitrile/water 84/16, v/v) for 1 hr. Extracts (4.0 mL) were filtered through a C18/alumina (1:3, w/w) column, and an aliquot (2.0 mL) of the supernatant was dried under compressed air at 50°C for an hour. The dried samples were derivatized with 100 µL of BSA: TMCS: TMSI (3:2:3, v:v:v). An aliquot of 1.0 mL of isooctane consisting Mirex (0.5 µg/mL) as an internal standard was added into the derivatized sample, and then the derivatization was terminated by adding the 1.0 mL of NaHCO3 (3%, w/v) solution. The derivated mycotoxins extracted in the isooctane layer were analyzed on an Agilent 6890N gas chromatograph with 5973 Mass Selective Detector (Agilent Technologies, Santa Clara, CA) under SIM mode. The separation was performed on a 35% phenyl siloxane column (30.0 m × 250 µm, 0.25 µm nominal) (Agilent HP-35, Part Number 19091G-133) as previously described. The ions of 295.20, 422.20, 512.30, 497.30 were used for the DON derivative qualification, 377.20, 392.20, 467.20, 482.30 for 3-ADON, 392.20, 407.20, 467.20, 482.20 for 15-ADON, 379.20, 482.30, 510.30, 585.30 for NIV, and 271.90, 236.90, 331.90, 403.70 for Mirex. The ion of 271.90 was used for the quantification of Mirex, and the areas of 295.20, 377.20, 392.20, and 271.90 were used to make the standard curves for quantifying DON, 3-ADON, 15-ADON, and NIV, respectively. The LOD and LOQ for these type B trichothecenes were 0.10 mg/kg and 0.20 mg/kg, respectively.

### CLSM examination of cereal grains and malt

Ten kernels of each sample of grain and malt were soaked in 10% neutral buffered formalin (10% NBF) solution for 12, 24, or 48 hrs at room temperature to fix and ease for sectioning. Fixed kernels were dehydrated in a series of alcohol washes and then 2-5 kernels were embedded in a paraffin wax block. Transverse sections (10 µm thickness) were taken from the middle portion of kernels within the paraffin block on a manual microtome (Leica, Heidelberg, Germany). Sections were collected on Superfrost plus charged slides (ThermoFisher Scientific, Waltham, MA, USA) for better attachemt (Solanki et al. 2020) and stained with 20 µg/ml of WGA-Alexa Fluor 488 (Invitrogen, Eugene, OR, USA) as described by Solanki et al. (2019). However, the staining time was optimized and extended to 5 hrs without shaking for better penetration of stain. The staining solution was replenished after 2.5 hrs to maintain the strength of fluorescent activity. Prepared slides were visualized on an LSM 700 laser microscope (Zeiss, Oberkochen, Germany) under a 40 × magnification oil immersion lens and 10 × objective lens using two different fluorescence channels. Green channel 488 was assigned for WGA-Alexa Fluor staining and red channel 555 was used for plant tissues with autofluorescence detection. To visualize inner structures in the section, Z-stack images were obtained (20-100 images per infection site) and were processed on Imaris 9.0.1 software for the 3D visualization of the fungal infection in the samples (Zeiss, Oberkochen, Germany).

### SEM examination of grain and malt kernels

Grain and malt kernels were cut in half transversely with a razor blade to expose the interior and attached to cylindrical aluminum mounts with colloidal silver paint (SPI Supplies, West Chester, Pennsylvania, USA). Then they were sputter-coated (Cressington 108auto, Ted Pella, Redding, California USA) with a conductive layer of gold. Images were obtained with a JEOL JSM-6490LV scanning electron microscope (JEOL USA, Inc., Peabody, Massachusetts USA) at an accelerating voltage of 15 kV.

## ACKNOWLEDGEMENTS

This material is based upon work supported by the U.S. Department of Agriculture under Agreement No. 59-0206-9-064, and also by the American Society of Brewing Chemists (ASBC Research Council). The USDA support is a cooperative project with the U.S. Wheat & Barley Scab Initiative. Any opinions, findings, conclusions, or recommendations expressed in this publication are those of the authors and do not necessarily reflect the view of the U.S. Department of Agriculture and ASBC.

This material is based upon work supported by the National Science Foundation under Grant No. 0619098. Any opinions, findings, and conclusions or recommendations expressed in this material are those of the author(s) and do not necessarily reflect the views of the National Science Foundation.

