## Supplementary material for "Localization of hyphal growth associated with mycotoxin production during the malting of Fusarium head blight infected grains": S1

### Slide 1
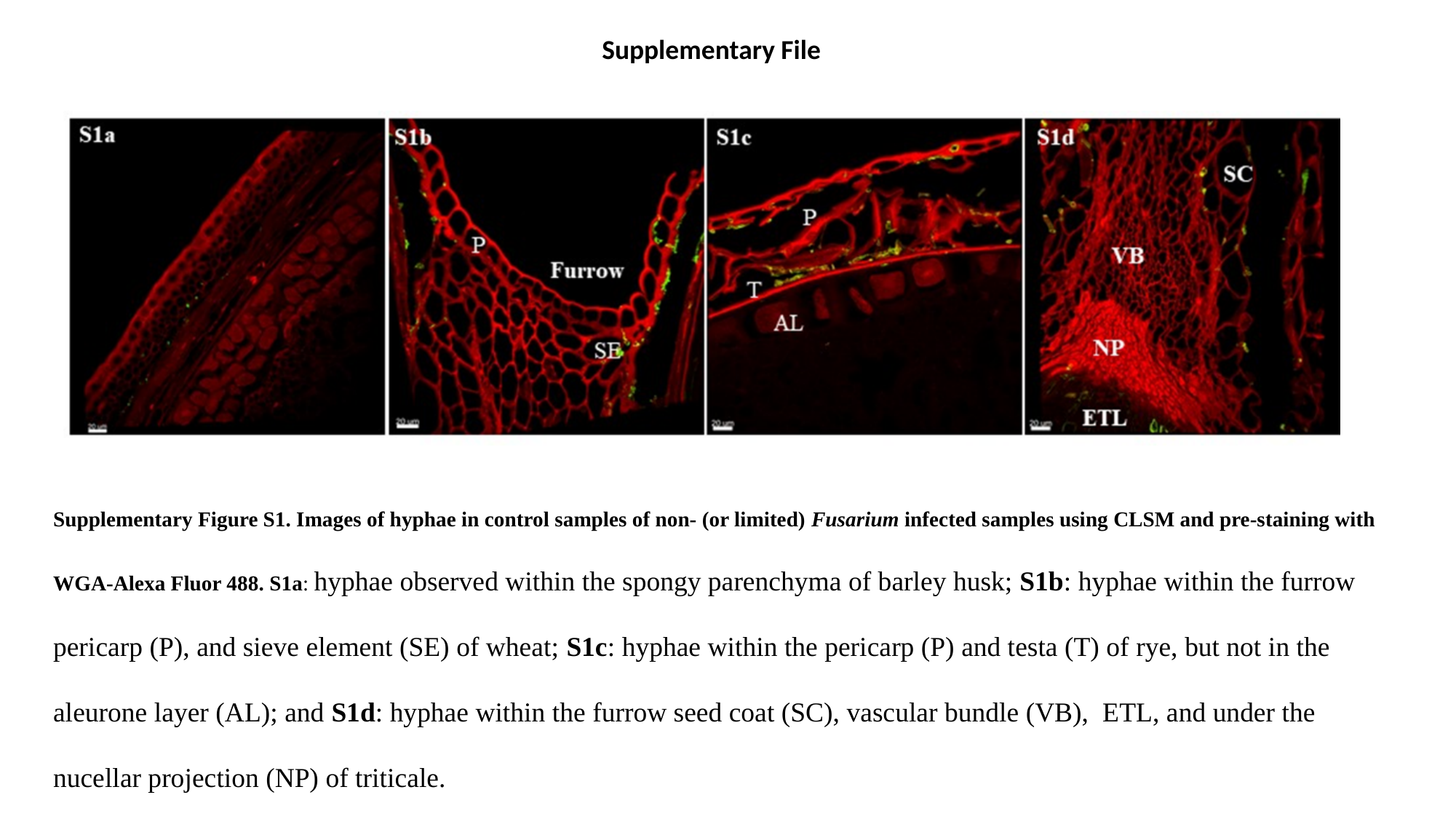

Supplementary File
Supplementary Figure S1. Images of hyphae in control samples of non- (or limited) Fusarium infected samples using CLSM and pre-staining with WGA-Alexa Fluor 488. S1a: hyphae observed within the spongy parenchyma of barley husk; S1b: hyphae within the furrow pericarp (P), and sieve element (SE) of wheat; S1c: hyphae within the pericarp (P) and testa (T) of rye, but not in the aleurone layer (AL); and S1d: hyphae within the furrow seed coat (SC), vascular bundle (VB), ETL, and under the nucellar projection (NP) of triticale.
